# Genetically Identical Mice Express Alternative Reproductive Tactics Depending on Social Conditions in the Field

**DOI:** 10.1101/2023.05.25.542282

**Authors:** Matthew N. Zipple, Caleb C. Vogt, Michael J Sheehan

**Author notes:** These authors contributed equally (Matthew N. Zipple, Caleb C. Vogt).

## Abstract

In many species, establishing and maintaining a territory is critical to survival and reproduction, and an animal’s ability to do so is strongly influenced by the presence and density of competitors. Here we manipulate social conditions to study the alternative reproductive tactics displayed by genetically identical, age-matched laboratory mice competing for territories under ecologically realistic social environmental conditions. We introduced adult males and females of the laboratory mouse strain (C57BL/6J) into a large, outdoor field enclosure containing defendable resource zones under one of two social conditions. We first created a low-density social environment, such that the number of available territories exceeded the number of males. After males established stable territories, we introduced a pulse of intruder males and observed the resulting defensive and invasive tactics employed. In response to this change in social environment, males with large territories invested more in patrolling but were less effective at excluding intruder males as compared to males with small territories. Intruding males failed to establish territories and displayed an alternative tactic featuring greater exploration as compared to genetically identical territorial males. Alternative tactics did not lead to equal reproductive success—males that acquired territories experienced greater survival and had greater access to females.

## Introduction

To deal with dynamic and unpredictable physical and social environmental conditions, animals are predicted to evolve plastic behavioral responses that allow them to make the best of a wide range of scenarios [1,2]. When different environmental conditions lead to different optimal reproductive behaviors, these plastic behaviors are referred to as “alternative reproductive tactics” or “conditional reproductive strategies” [3,4]. For many species, establishing and maintaining a territory is a central aspect of individuals’ reproductive life history, as territorial control allows them to reliably access physical resources and attract mates [5–11]. We therefore expect behaviors related to territory formation, defense, and invasion to have been under strong selection in these species and for animals to plastically alter their territorial behavior in response to a wide range of social environmental conditions.

Animals seeking to establish territories may encounter radically different social environments that vary widely in their intensity of competition. At one extreme, animals may seek to establish a territory in a relatively unoccupied environment with an abundance of resources and a lack of competitors for space. This is the situation faced by, for example, rodents living in low-density populations at the start of a breeding season [12–15] or by the earliest migratory birds to arrive at a breeding ground [16–20]. On the other extreme, an animal might develop or compete in a world where suitable territories are either largely or entirely filled. Such is the world often encountered by rodents born later into a breeding season after colonization and population growth has already occurred or migratory birds arriving relatively late to a breeding ground [12–20].

If the exact same animal found itself in a more or less competitive social environment, would its territorial behavior look different? How so? In many species, males who are unable to establish territoriality control or social dominance adopt an alternative “sneaker” tactic to attempt to furtively mate with females as a conditional strategy to make the best of a bad situation [4,21–25]. Yet, in natural populations it is difficult to know whether these differences in tactics are caused by an individual’s quality, its history of social interactions, or the broader current social context in which it lives. The simplest way to establish unambiguous causality regarding the effect of social environment on individual behavioral decisions is by manipulating a single aspect of social environment while holding genotype and developmental conditions constant. But such manipulations of environmental conditions are rarely, if ever, possible in wild populations [26].

Experimental populations of inbred mouse strains (*Mus musculus domesticus*) living in seminatural enclosures provide the ideal opportunity for studying the causal impact of social environment on individual competitive and reproductive behaviors. Wild and lab mice establish and defend territories when given the space to do so, and territories allow males to monopolize or nearly monopolize access to food and mates [27–35]. And the identical genetics and standardized rearing conditions of inbred strains represent an extreme uniformity across individuals as compared to wild populations, allowing us to manipulate a single aspect of animals’ social environments and draw causal conclusions about the impact of this manipulation [26].

In this paper, we characterize the behavioral tactics of genetically identical mice that either encounter (a) a world of abundant, unfilled territorial spaces and limited conspecific competition or (b) a world in which residents already occupy territories. The resulting data allow us to test the hypothesis that animals with similar prior experiences will rapidly develop alternative tactics in response to the current social environment in which they find themselves. Additionally, we use this data to test three hypotheses regarding mouse territorial behavior, in particular: (1) that territory size is constrained by social factors, such that males with larger territories face greater invasion pressure than males with smaller territories, (2) that territorial males monitor their social environment and respond to salient changes in it and (3) that territories confer benefits to males in the form of both survival and access to females. The data also allow us to describe the dynamics of territory formation and defense in the most studied biomedical model organism in finer-grain detail than ever before. Given recent public attention to the constraints of the laboratory environment on drawing useful inferences from lab mice, this latter contribution is particularly timely [36].

## Materials and Methods

### Field Enclosure and Study Subjects

A detailed description of the enclosures at Cornell University’s Liddell Field Station can be found elsewhere [37], so here we will only describe those elements critical to the success of this experiment. The enclosure is 15m x 38m, approximately 9,000 times the area of a typical mouse cage. Within the enclosure we set up 12 plastic tubs (31 gallon storage totes, Rubbermaid, USA), placed in an equally-spaced 3×4 grid across the enclosure (**Figure S1**). Each tub (hereafter “resource zones”) contained *ad libitum* food access and provided insulation and shelter from adverse weather conditions. We equipped each zone with a single entrance and exit made out of a 6-inch-long PVC pipe (2” in diameter). These resources and the single entrance made the resource zones highly valuable, defendable areas that are meant to mimic the foraging landscape of commensal mice. To track the comings and goings of mouse visitors to each zone, we placed a 10-inch RFID antennae (Biomark, USA), beneath the entrance tube of each zone. The antennas were connected to a central monitoring system (Small Scale System, Biomark, USA) and transmitted RFID reads at a rate of 2-3 Hz.

Our study subjects were 20 male and 20 female eight-week-old lab mice (C57BL/J6 strain), obtained from The Jackson Laboratory. After arrival at our lab, we separated individuals into smaller holding cages containing either 2 males or 4 females. After allowing animals to acclimate for 8 days, we administered isoflurane (an inhaled anesthetic) and injected two subcutaneous passive integrative transponders (PIT) tags in the flank and between the scapulae of each mouse (MINI HPT10, Biomark, USA) using 16-gauge needles. Inserting two PIT tags allowed us to continue to monitor individuals in the field even if one of the tags was lost. Based on past experience, we anticipated PIT tag loss at < 5%, making it quite unlikely that any individual mouse would lose both tags during the experiment.

### Manipulating the Social Environment of Genetically Identical Animals

On the afternoon of September 24, 2021 we simultaneously released 8 male and 8 female mice in the center of the enclosure. We allowed mice to explore the enclosure and establish territories over the first five nights of the experiment. During this initial stage the number of male mice (8 animals) was substantially smaller than the number of resource zones (12 zones). These animals were entering a world of abundant resources with relatively few competitors.

Then, on the afternoon of September 29 (the 6^th^ night of the experiment) we released 12 additional males (hereafter ‘intruding’ males) and 12 additional females into the enclosure. We observed mouse movement and spatiotemporal dynamics between territorial and intruding males for the next two weeks, after which point a substantial number of intruding males appeared to have died (they no longer visited any zones despite having visited previously and were never captured during subsequent trapping efforts). We then allowed the population to persist for an additional 15 days (35 days total from the beginning of the experiment) to continue to measure differential survival outcomes between territorial and intruder males before trapping and removing all surviving animals.

### RFID Data Analysis

For all analyses below, we used the data collected from the RFID system. We calculated the number of zones that animals visited each night to assess the breadth of animals’ movement in the enclosures. We also identified movements between zones each time that an animal appeared in one zone followed by appearing in a different zone. To assess territorial control, we calculated the proportion of male-sourced reads at a zone originating from the male with the highest proportion of reads on each night.

For social network analyses, we inferred the amount of time that animals overlapped in the same zone based on their patterns of RFID reads. We have described the process for inferring the duration of overlap elsewhere in detail [37]. Briefly, if a mouse registered consecutive RFID reads in the same zone within a given time window, we assume that the mouse had been in the zone for the period between those reads. Because the zones are ∼400% larger than the area of the antenna, mice will often spend substantial time in a zone but only register RFID reads occasionally. To estimate the duration of different visits to a given zone we first identified the 95^th^ percentile for the amount of time that passed between reads of the same individual in the same zone across all individuals and all zones in our experiment (211 seconds). If a mouse registered an RFID read in the same zone with less than this length of time passing between reads, we assumed that it had been present in the zone for the entirety of the interim period. We then calculated periods of spatiotemporal overlap with other animals. While this assumption about animals’ presence in the zone is of course imperfect, this approach provides a noisy but informative view of the social world of these animals.

### Statistical Analyses

We performed all statistical analyses in R. We built mixed effects models using the glmmTMB package [38]. For each analysis any transformations of response or predictor variables were chosen based on visual inspection of the relationship between the two variables as well as the resulting residuals from models of untransformed variables. We included relevant random intercepts and random slopes in each mixed effects model, as appropriate. We identify the random effects structure for each analysis in the results tables below. We performed the repeatability analysis described below using the rptR package [39] and the time-varying survival analysis using the survival package [40].

## Results

In this experiment, we exposed genetically identical, age-matched male mice to two different social environments—one in which territories were empty and resources were abundant and one in which territories were full and resources were restricted. Below we first describe the social and spatial behavior of the first group of males in an empty social environment, followed by their different reactions to the addition of the second group of males. We then compare the alternative socio-spatial behavior of the two groups of males, depending on the social environment that they encountered. We close by describing the differential survival and apparent reproductive outcomes obtained by the males that encountered the two different social environments.

### Behavior of Males Entering an Empty Social Environment

For the first five days in the enclosure, the eight original males experienced an environment with abundant resources and relatively low levels of competition. During this time, the number of available resource zones exceeded the number of males, and the eight males rapidly established territorial control over each of the twelve resource zones. Across all 12 zones, the proportion of all RFID reads belonging to the eventual territory male increased during the first five nights of the experiment, such that nearly all (99.97%) of those reads recorded on night 5 were reads from the territory holder (**Figure S2**). The pattern of increasing control over each zone by a single male resembles previous patterns observed for this strain in a previous experiment [37].

By night 5, each male accounted for the majority of male reads in either one (n = 4) or two (n =4) resource zones. Males displayed strikingly different patterns of space use depending on the number of zones contained within the territories that they established. Those males that established territories containing a single zone (hereafter “one-zone males”) very rarely visited another zone (**Figure 1**), averaging only 2.5 transitions between zones each night during these first five nights of the experiment. In contrast, males holding two territories (hereafter “two-zone males”) consistently spent time in one resource zone during the day and made frequent excursions between the two zones at night (**Figures 1, S3**), averaging 11.0 transitions between zones during the same period.

**Figure 1.**
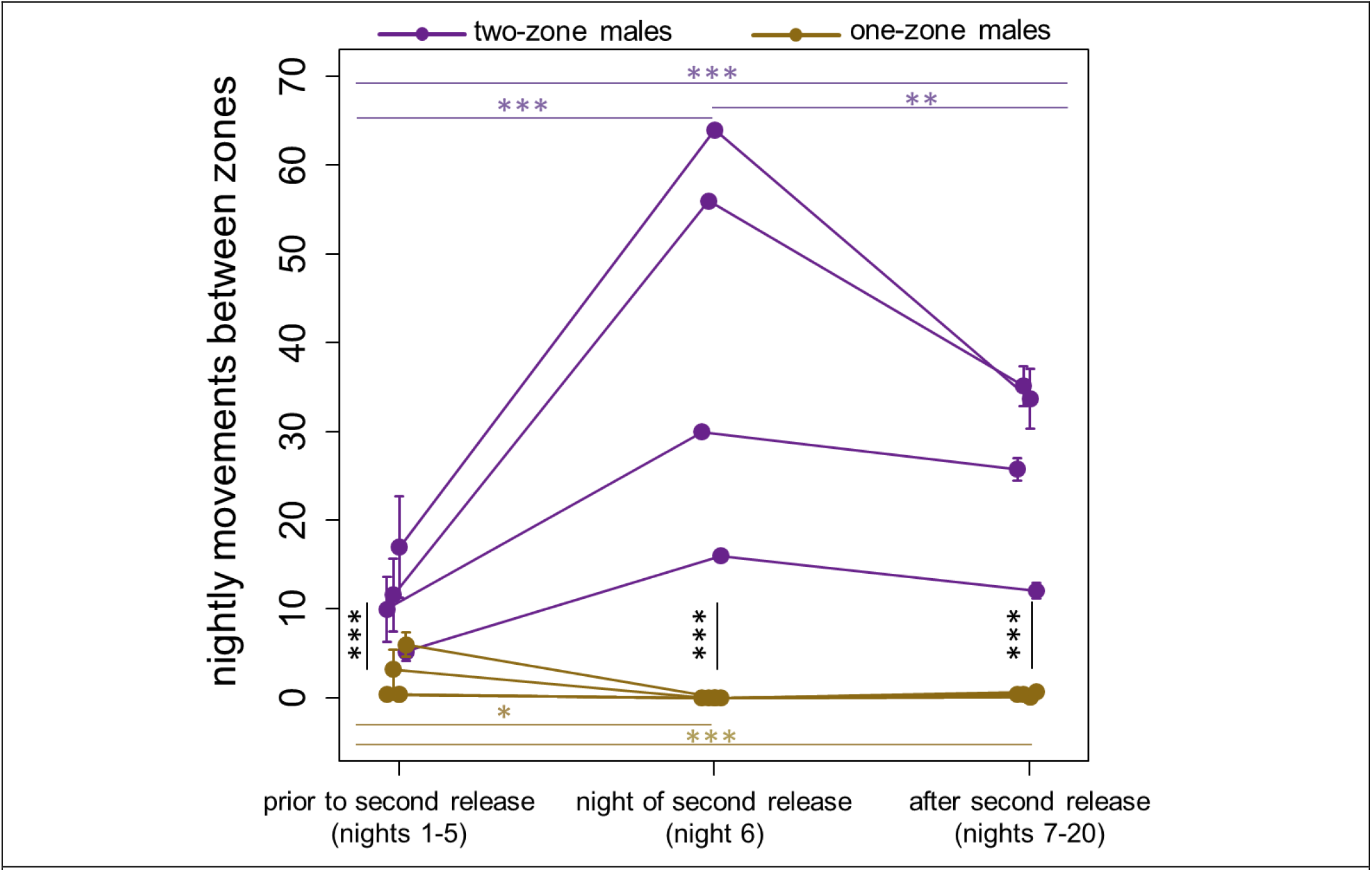
Males responded to the introduction of intruder males differently, depending on the size of their territory. The y-axis represents the average number of nightly transitions between resource zones that males performed, with each point representing a single male during a different period of the experiment. Males with larger territories, containing two resource zones (purple points and lines) increased their number of nightly trips between zones in response to the introduction of intruding males on night 6, and maintained this elevated patrolling behavior thereafter. In contrast, males with smaller territories, containing only one resource zone (gold points and lines) responded by reducing their number of nightly transitions between zones and essentially never moved between zones again. Asterisks indicate levels of statistical significance for comparisons, extracted from mixed-effects models (* p < 0.05, ** p < 0.01, *** p < 0.001; Table 1).

#### Territory size influences resident male behavior in the face of intruders

On day 6 of the experiment, we added an additional 12 males (hereafter ‘intruder males’) and 12 females to the enclosure. Territorial males responded differently to this introduction depending on whether they held one or two resource zones within their territory. On the night of the introduction, two-zone males responded by significantly increasing the frequency with which they moved between their two zones (p < 0.0001). The magnitude of this increase varied among these four males, but was substantial in all four cases, ranging from a 200% to 383% increase as compared to the average number of zone transitions during their first five nights (**Figure 1**). In contrast, males holding a territory containing a single resource zone significantly *decreased* the number of nightly transitions that they made between zones—these males essentially never moved between zones again after the introduction of additional males (**Table 1, Figure 1**). These results indicate (1) that males were monitoring their social environment and scaling their behavior in response to changes in it, (2) that males with larger territories needed to expend more energy on patrolling and defending their territories as compared to males with smaller territories, and (3) that this energetic cost of territory size was especially acute under dense social conditions, when intruder males were present (i.e., after night 5 of the experiment).

**Table 1.**
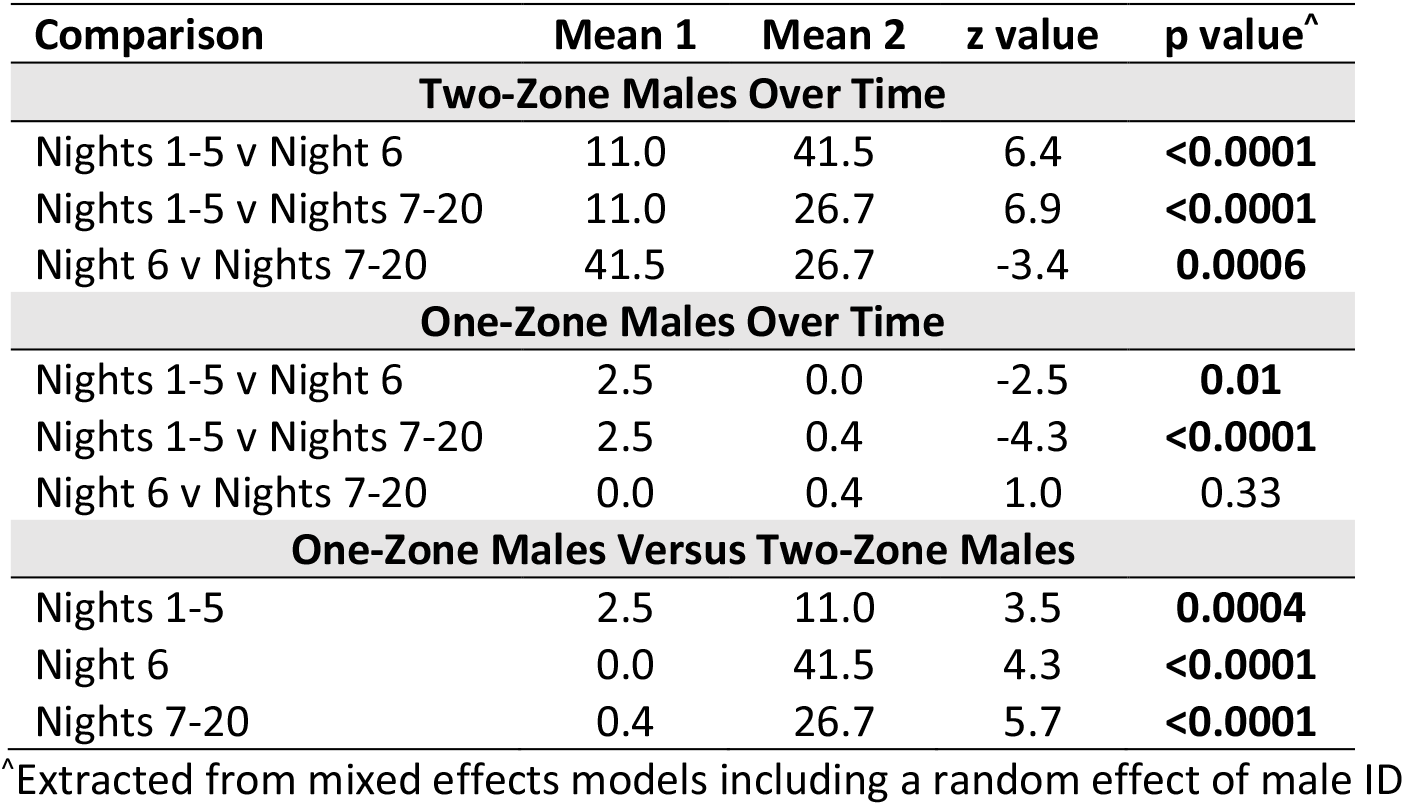
Comparisons of the average number of transitions between zones made by males with different territory sizes at different points in the experiment.

| Comparison | Mean 1 | Mean 2 | z value | p value<sup>^</sup> |
| --- | --- | --- | --- | --- |
| **Two-Zone Males Over Time** |  |  |  |  |
| Nights 1-5 v Night 6 | 11.0 | 41.5 | 6.4 | **<0.0001** |
| Nights 1-5 v Nights 7-20 | 11.0 | 26.7 | 6.9 | **<0.0001** |
| Night 6 v Nights 7-20 | 41.5 | 26.7 | -3.4 | **0.0006** |
| **One-Zone Males Over Time** |  |  |  |  |
| Nights 1-5 v Night 6 | 2.5 | 0.0 | -2.5 | **0.01** |
| Nights 1-5 v Nights 7-20 | 2.5 | 0.4 | -4.3 | **<0.0001** |
| Night 6 v Nights 7-20 | 0.0 | 0.4 | 1.0 | 0.33 |
| **One-Zone Males Versus Two-Zone Males** |  |  |  |  |
| Nights 1-5 | 2.5 | 11.0 | 3.5 | **0.0004** |
| Night 6 | 0.0 | 41.5 | 4.3 | **<0.0001** |
| Nights 7-20 | 0.4 | 26.7 | 5.7 | **<0.0001** |

<sup>^</sup>Extracted from mixed effects models including a random effect of male ID

No successful takeover event appeared to occur during the two weeks following the introduction of new males (through night 20 of the experiment). A successful takeover would have been visible in our data as an event in which a new male became responsible for a plurality of RFID reads within a zone on a given night and maintained that position thereafter. In two cases, an intruder male was responsible for a plurality of RFID reads at an antenna for a brief period, but the original territorial male then quickly reclaimed the territory.

#### Relative defensibility of differently sized territories

Given their increased effort to maintain the integrity their territories, we next asked whether two-zone males were able to defend their territories with a comparable degree of success as one-zone males. **Figure 2** displays the average proportion of reads in a given zone that originated from the territory-holding male, depending on whether that male held one or two zones in his territory. Although there was no significant difference between these values on night 5 of the experiment (p = 0.45, before the introduction of new males), a large difference emerged following the introduction of additional males on night 6.

**Figure 2.**
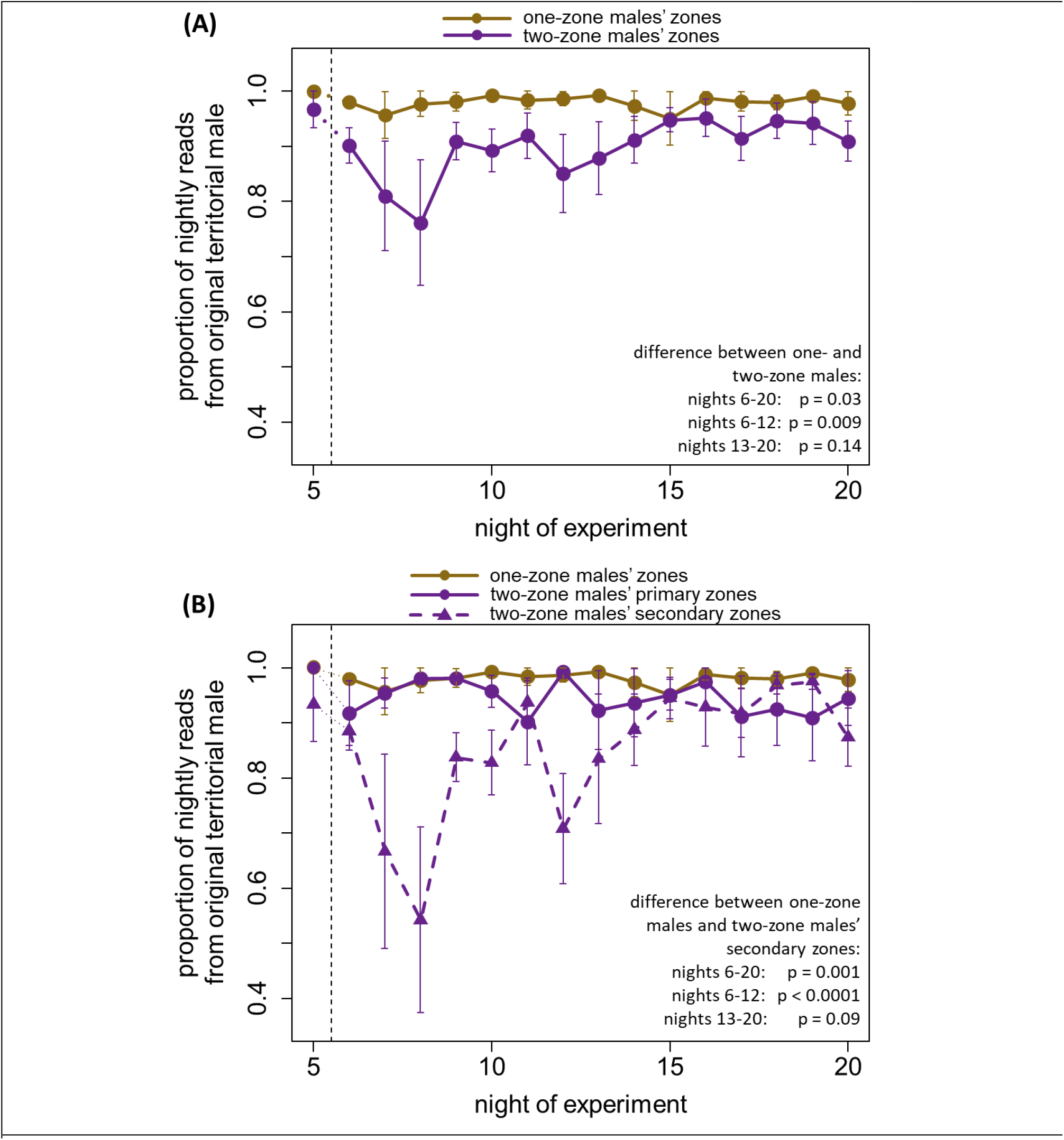
Two-zone males were somewhat less able to defend their territories from intruders. (A) The y-axis represents the average across zones of the proportion of nightly RFID reads that originated from the territorial male that controlled the zone. Higher values correspond to a zone being more defensible and suffering fewer incursions by non-territory holders. Following the introduction of new males (indicated by the vertical dashed line), zones contained in larger territories became significantly less defendable than zones contained in smaller territories. (B) This difference in defensibility was true only of one of the two-zone males’ zones (their ‘secondary’ zones). Two-zone males were able to maintain territorial integrity comparable to one-zone males in their primary zones. In both panels, p values refer to mixed effects models that included random effects of territorial male ID.

While one-zone males experienced only a negligible reduction in their ability to exclude other males from their territories, zones controlled by two-zone males experienced substantial incursion (**Figure 2A, Table 2**). Across nights 6 through 20, the proportion of reads in a given zone belonging to the territory holder was significantly lower if the territory-holder was a two-zone male (mean = 0.90) rather than a one-zone male (mean = 0.98, difference: p = 0.03). This effect was strongest during the week starting on the night of male introduction (nights 6-12), when the mean proportion of reads from the territory-holder was only 0.86 in zones held by two-zone males, but remained at 0.98 in zones controlled by one-zone males (p = 0.009).

**Table 2.** Comparisons of the average proportion of RFID reads in a given resource zone that originated from the territory holder (a measure of defensibility), depending on territory size.

| Period of Comparison | One-Zone Males' Zones | Two-Zone Males, Both Zones<br>(z value; p value)* | Two-Zone Males' Primary Zones Only<br>(z value; p value)* | Two-Zone Males' Secondary Zones Only<br>(z value; p value)* |
| --- | --- | --- | --- | --- |
| Night 5 | 1.00 | 0.97<br>(-0.8; 0.45) | 1.00<br>(0.0; 1.00) | 0.94<br>(-1.4, 0.16) |
| Nights 6-20 | 0.98 | **0.90**<br>**(-2.2; 0.03)** | 0.94<br>(-0.9, 0.37) | **0.85**<br>**(-3.3, 0.001)** |
| Nights 6-12 | 0.98 | **0.86**<br>**(-2.6, 0.009)** | 0.95<br>(-0.5, 0.59) | **0.77**<br>**(-4.5, <0.0001)** |
| Nights 13-20 | 0.98 | 0.93<br>(-1.5, 0.14) | 0.93<br>(-1.2, 0.24) | 0.92<br>(-1.7, 0.10) |

^Extracted from mixed effects models including a random effect of territory holder ID
\*All z and p values are in comparison to one-zone males' zones

Additional investigation revealed that two-zone males did not suffer incursions into their two zones at equal rates. Instead, two-zone males appeared to prioritize defensive attention on one of their two zones, from which they were able to almost entirely exclude intruding males (their “primary” zone, **Figure 2B, Table 2**), mirroring the ability of one-zone males. In contrast, the second zone that they controlled (their “secondary” zone) was significantly less defendable than zones controlled by single males (**Figure 2B, Table 2**).

### Behavior of Males Entering a Filled Social Environment

The males that we added on night 6 of the experiment entered a filled social environment that lacked any available resource zones. Although no intruding males successfully took over any resource zones in the first two weeks after their addition, they did make frequent visits to existing males’ territories. Intruding males’ space use and exploratory behavior changed over the course of the experiment. Intruding males explored the greatest number of zones on their first night in the enclosure (mean = 2.9, 95% CI: 2.2-3.7), before visiting fewer zones on ensuing nights (mean = 1.8, 95% CI: 1.4-2.1, p = 0.0004, **Figure 3A, Table 3**).

**Table 3.** Results from mixed effects models regarding the behavior of intruding males.

| Parameter | Estimate | Std. Error | z value | p value | Interpretation |
| --- | --- | --- | --- | --- | --- |
| Number of Zones That Intruding Males Visited on a Given Night<sup>1</sup> |  |  |  |  |  |
| Intercept (Reference = Night 6) | 2.9 | 0.3 |  |  | Intruding males visited more zones on their first night in the enclosure as compared to subsequent nights. |
| Nights 7-12 | -1.2 | 0.3 | -3.5 | **0.0004** |  |
| Nights 13-20 | -1.1 | 0.3 | -3.3 | **0.001** |  |
| Probability That an Intruding Male Visited a Given Zone on a Given Night<sup>2</sup> |  |  |  |  |  |
| Intercept | -3.3 | 0.5 |  |  |  |
| Zone Held by Two-Zone Male | 0.7 | 0.7 | 0.9 | 0.34 |  |
| Did the Same Male Visit the Zone on the Previous Night? (yes vs no) | 1.3 | 0.4 | 3.4 | **0.0007** | An intruding male was much more likely to visit a zone if he had visited the same zone the previous night. |
| Zone Held by Two-Zone Male x Same Male Visited Yesterday | 1.2 | 0.4 | 2.7 | **0.006** | The effect of visiting a zone on the previous night was stronger when the zone was controlled by a two-zone male |

<sup>1</sup>Estimates extracted from linear mixed models including random effects of intruding male ID248 <sup>2</sup>Estimates extracted from generalized linear mixed model (binomial error distribution) including random effects of intruding male ID, territorial male ID, and zone ID as well as a random slope of territory size nested within intruding male ID.

**Figure 3.**
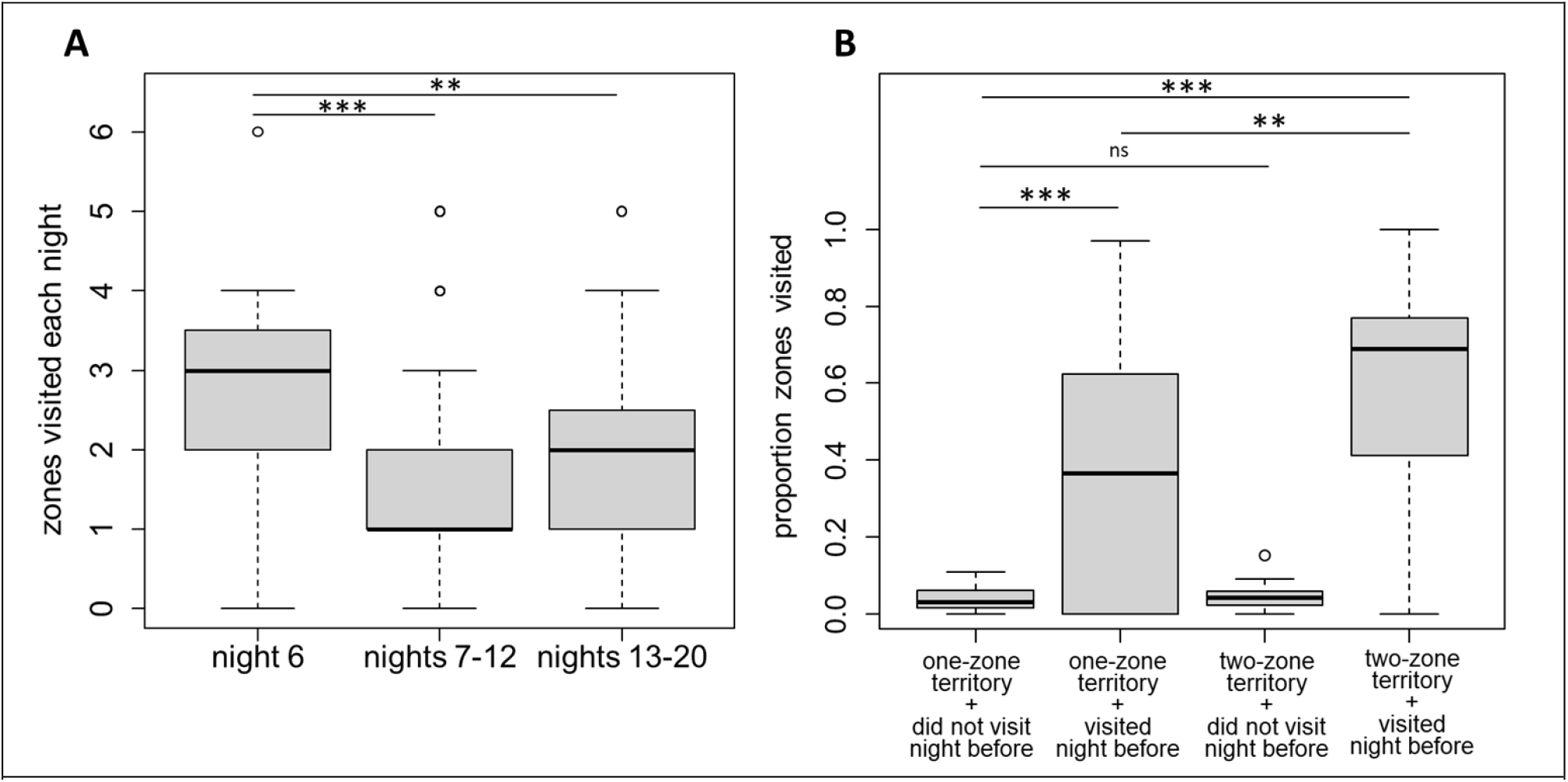
The zone visitation patterns of intruding males. (A) Intruding males made nightly visits to occupied resource zones, but visited fewer zones after their first night in the enclosure (night 6). Boxplots represent the distribution of nightly visits in each period for all intruding males. (B) Intruding males were more likely to visit a zone on a given night if they had visited the zone the night before. This site fidelity was especially strong when the zone belonged to a two-zone male. In both panels significant differences (identified with a mixed effects model) are identified with asterisks (** p < 0 .01, *** p < 0.001)

Given our finding that zones controlled by two-zone males were more prone to intrusion by non-territorial males (**Figure 2, Table 2**, above), we sought to gain a better understanding of the decision-making processes among intruding males that led to this outcome. To do so, we built a mixed effects logistic regression model to interrogate the decision-making process at the level of an individual intruding male mouse on a given night. This analysis yielded two results. First, intruding males appeared to show some spatial fidelity, despite not holding territories in resource zones. Intruding males were much more likely to visit a zone on a given night if they had visited that zone on the previous night (p < 0.0001, see **Figure 3B, Table 3**). And this site fidelity was especially strong when the zone the intruder had visited the night before belonged to a two-zone male (interaction with territory size, p = 0.006, **Figure 3B, Table 3**).

### Alternative Tactics Between Males Entering Different Social Environments

We found strikingly different patterns of exploratory behavior, depending on whether males entered an environment full of unoccupied territories (the first males) or instead entered an environment in which all territories were already occupied (**Figure 4**). While both sets of males explored similar numbers of resource zones during their first three nights in the enclosure, the original males who were able to find and acquire territories largely ceased exploration after these first three nights. In fact, after these first three nights, 4 of the 8 original males never entered a new zone again during their next 12 nights in the enclosure (the remaining 4 entered 1 or 2 new zones each, mean for all 8 = 0.6,see **Figure S4**). In contrast, the males added on night 6 (who were unable to establish territories in the zones that they had explored after 3 nights) continued to explore new zones (mean new zones among surviving intruders = 4.2 zones, interaction between status and time in enclosure, p = 0.0007, Table 4). The outcome of this difference in exploratory behavior was that the group of intruding males on average had explored substantially more zones by their 15^th^ night in the enclosure than the original territorial males (night 20 of the experiment, 7.5 zones versus 5.7 zones, **Figure 4**). A comparable analysis that considers as the unit of analysis the number of nightly new zones that a male visited yields the same results (see **Figure S4**).

**Table 4.** Results from a mixed effects model predicting the number of cumulative unique zones visited by a male after their first three nights in the enclosure.

| Parameter<sup>1</sup> | Estimate | Std error | z value | p value | Interpretation |
| --- | --- | --- | --- | --- | --- |
| Intercept<br>(Reference = Original Males) | 5.3 | 0.8 |  |  |  |
| Total Nights Spent in<br>Enclosure (nights 3-15) | 0.03 | 0.05 | 0.6 | 0.54 | Original males visited very few new zones after their first three nights in the enclosure |
| Male was an Intruding Male | -0.9 | 1.0 | -0.9 | 0.36 | Original and intruding males visited a comparable number of unique zones during their first three nights in the enclosure. |
| Intruding Male x Nights | 0.23 | 0.07 | 3.4 | **0.0007** | Intruding males continued to visit new zones throughout their time in the enclosure |

<sup>1</sup>Results are from a linear mixed effects model that also included a random intercept of male ID along with a random slope of total nights spent in the enclosure

**Figure 4.**
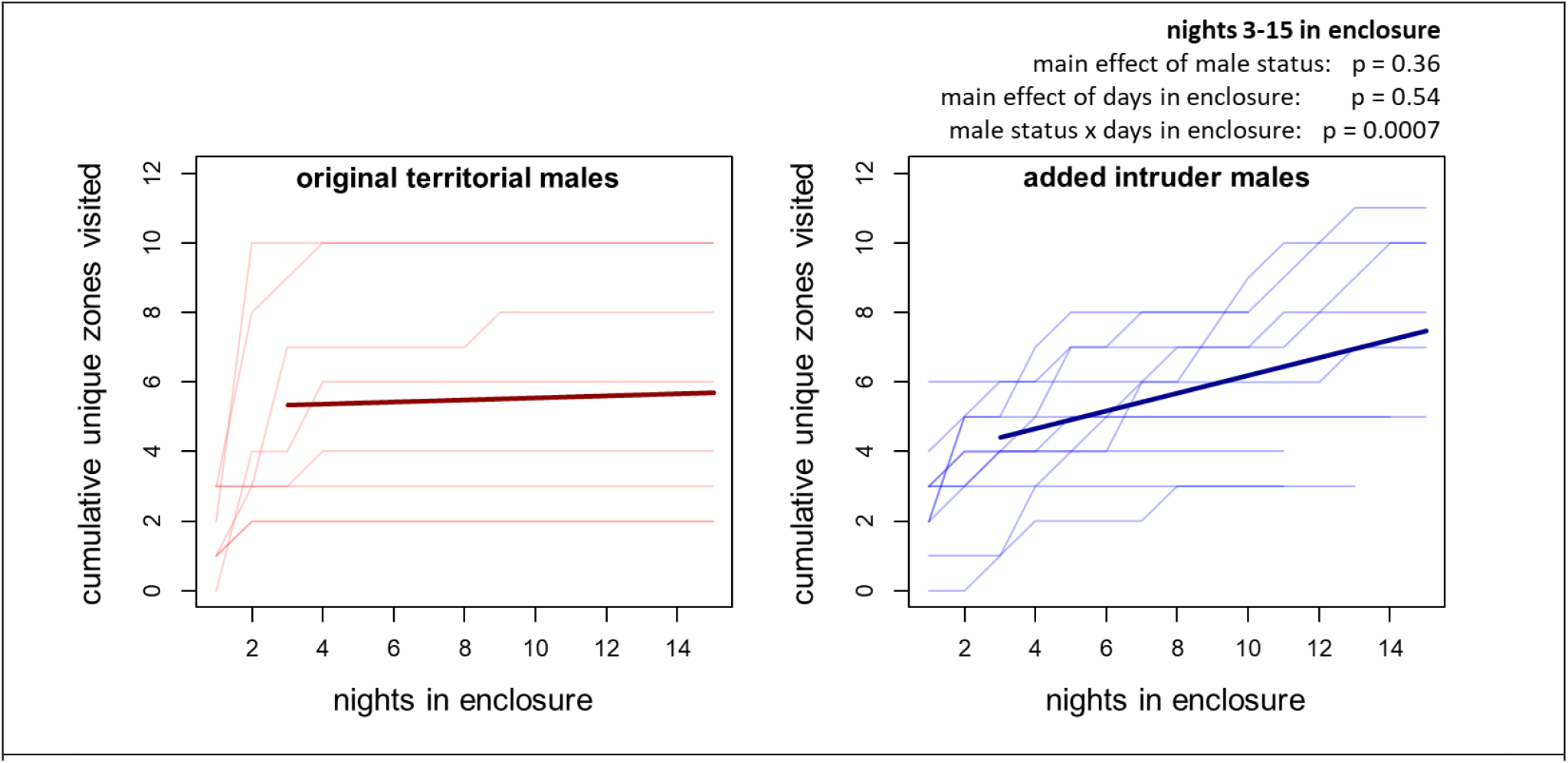
Male exploratory behavior differed, depending on whether the male encountered an environment without any occupied territories (left) or an environment with all territories already filled (right). The y-axis represents the cumulative number of resource zones that males visited and the x-axis represents how many days males had been in an enclosure. Intruding, but not territorial, males continued to explore the enclosure after initial exploration, such that the intruding males had visited substantially more zones by the end of the experiment. Faint lines represent data from individual males and thick curves represent model estimates from the mixed effects model described in Table 4.

A cursory examination of **Figures 3 and 4** reveal substantial variation in intruding males’ space use, which may reflect differences in efforts to explore and monitor territories and the males that controlled them. Indeed, while some intruding males generally visited one zone each night after their first night in the enclosure, others consistently visited 2 or more zones. Overall, male identity explained an estimated 25% of the variance in the number of zones that an intruding male visited each of nights 7-20 of the experiment (95% conf. int. = [0.06,0.45], p < 0.0001). In the current paper we are unable to assess whether such variation in space use among intruding males eventually shapes eventual territory acquisition or reproductive success, but the presence of such variation suggests a fruitful path for future studies.

### Survival and Reproductive Opportunities of Males Expressing Alternative Behavioral Tactics

Finally, we assessed whether males’ expressing alternative behavioral tactics achieved apparent differences in fitness, as measured by (a) survival and (b) access to females.

To assess the long-term survival dynamics in our enclosures, we allowed the experiment to continue for an additional 15 days after the end of our focused investigation of territorial defense dynamics (35 total days from the first introduction of our original males). Over these 35 days (**Figure 5**), males without territories died at significantly higher rates than did either (a) males with territories (hazard ratio = 5.9, 95% CI = 1.2-29.1, p = 0.03) or (b) females (hazard ratio = 4.7, 95% CI = 1.6-14.3, p = 0.005). Given the low levels of mortality in territorial males, we were unable to assess whether territory size (i.e. two zones versus one zone) had an additional effect on territorial male mortality.

**Figure 5.**
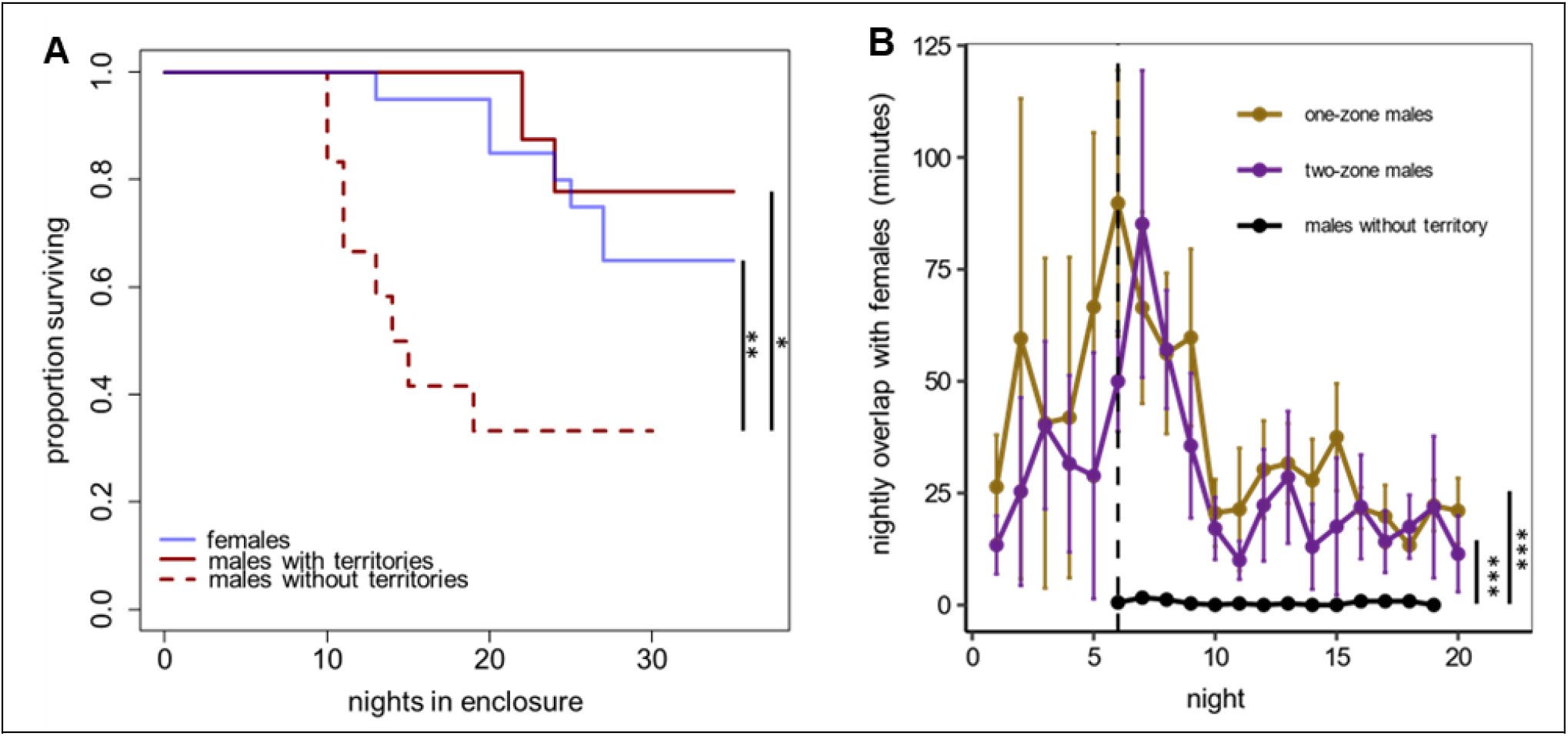
**(A)** Males with territories experienced a survival advantage as compared to males without territories and survived at comparable rates as females. **(B)** Males with territories spent more time overlapping with females in resource zones than males without territories. Asterisks indicate statistically significant differences between groups (*** p < 0.001,** p < 0.01, * p < 0.05).

We also assessed whether territorial males achieved greater access to females than males without territories. Territorial males spend much more time overlapping with females in resource zones than do males without territories (**Figure 5B**, lmm t_98_ > 4.3, p < 0.0001). independently replicating recently published results from a different study in this system [40]. Second, we find that one and two-zone males spend comparable amounts of time overlapping with females (**Figure 5B**, p > 0.05). Thus, we identify major differences in reproductive success in our system that were not the result of any differences in genetics or developmental environment, but instead were the causal result of the competitive social environment that a male happened to encounter.

## Discussion

By manipulating the social environment experienced by genetically identical, age-matched mice, we have identified causal impacts of the current social environment on individuals’ behavioral tactics. When placed in semi-natural field enclosures that reproduce ecologically relevant physical and social conditions, the canonical strain of lab mouse (C57BL/6J) expresses at least 3 alternative reproductive tactics depending on the present social environment that individuals encounter. Males that entered a world of abundant resources and a low number of competitors established territories and rarely left these spaces after establishing control over them. Their ability to control these spaces In contrast, age-matched, genetically identical males that encountered a filled social landscape without available territories failed to monopolize space and instead continued to explore a wider range of the physical space in the enclosure. Within territorial males, the size of their territory and the addition of intruder males had strong impacts on their space use and movement patterns, indicating that males are acutely aware of changes to their social environment and alter their behavior in response to such changes.

Unlike many studies of conditional strategies or alternative mating strategies under natural conditions, which examine the role of nutritional [41,42], abiotic [43], or genetic [44,45] factors in determining behavior, here we controlled genetic and developmental variation by using a single inbred strain of lab mice. The only difference between those males expressing territorial behavior and those expressing intruder behavior was the social environment into which they were placed. This study joins recent advances in lab-based manipulations of social status in monkeys [46] and mice [47] and social experience in flies [48] that reveal the individual and society-level impacts of variation in a controlled social environment. Though not measured here, our social manipulation likely also led to differences in males’ physiology and resource deployment, such as differences in gene expression or metabolism and signal allocation. In the lab, dominant animals show different gene expression profiles than subordinate animals [49] and animals that experience competitive success or failure rapidly alter their patterns of chemical signaling through urination [50].

The alternative reproductive tactics that males expressed were accompanied by apparent differences in survival and access to females. Males that entered a filled social environment and were forced to pursue a territory-less tactic died more quickly and spent less observed time overlapping with females while they were alive. Within the group of territory-holding males, maintaining larger territories appeared to come with a socially-imposed cost. After the addition of intruder males, those zones that were controlled by two-zone males were more vulnerable to incursion. Territories (in particular, ‘secondary zones’) were less well monopolized, and intruding males’ tendency to return to the same zones on subsequent nights was particularly strong when that zone was controlled by a two-zone male (**Figures 2 and 3**). This latter finding suggests that by visiting the territory of a two-zone male, intruders assess that the territory may be relatively porous or otherwise attractive, causing them to be particularly likely to return the following night (a version of a “win-stay, lose-shift” tactic [51]).

The primary limitation of this study is that we were only able to measure space use within the resource zones that we set up, which likely represent a small, though extremely important, part of a male’s territory. We infer that all territorial mice spent a substantial, but minority, portion of their daily in and immediately around these zones (on the order of 3-10 hours per day on average). We suspect that the remainder of males’ time was spent outside of zones, but within the rest of their territories, which we suspect comprised ∼10-30 square meters surrounding the zone(s) that the male controlled, as well as the series of tunnels that mice regularly dug below their zones. Still, we expect our measures of male space use within zones to largely predict space use within the larger territories outside of the zones. This expectation is bolstered by results from Smith et al [52], who report that in California ground squirrels space use below ground (inferred by a similar RFID approach taken here), strongly predicted above-ground social networks that were observed directly.

The approach that we take here—studying the impacts of variation in social environment in model organisms living outside of a highly artificial laboratory environment—holds great potential for additional advances [26]. By focusing on what is important to these animals’ natural history, in combination with using high-throughput approaches to study animals’ whose genetics, demography, and social environment we can control, we are able to test hypotheses and draw causal conclusions about behavior, individuality, and society. These same conclusions are extremely difficult if not impossible to make with unambiguous causality in either fully wild populations or the overly constrained social conditions of the lab.

## Acknowledgments

We gratefully acknowledge our sources of funding that made this work possible. MNZ has been supported by an NSF postdoctoral fellowship in biology (award # 2109636) and a Klarman postdoctoral research fellowship from Cornell University. CCV is supported by a Mong Neurotechnology Fellowship from Cornell University. This work was also supported by Pilot and Feasibility awards to MNZ and MJS from the Animal Models for the Social Dimensions of Health and Aging Network (project #5R24AG065172-03). The costs of care for the mouse colony were supported in part by R35 GM138284 to Andrew Moeller.

## Supplementary Figures

**Figure S1.**
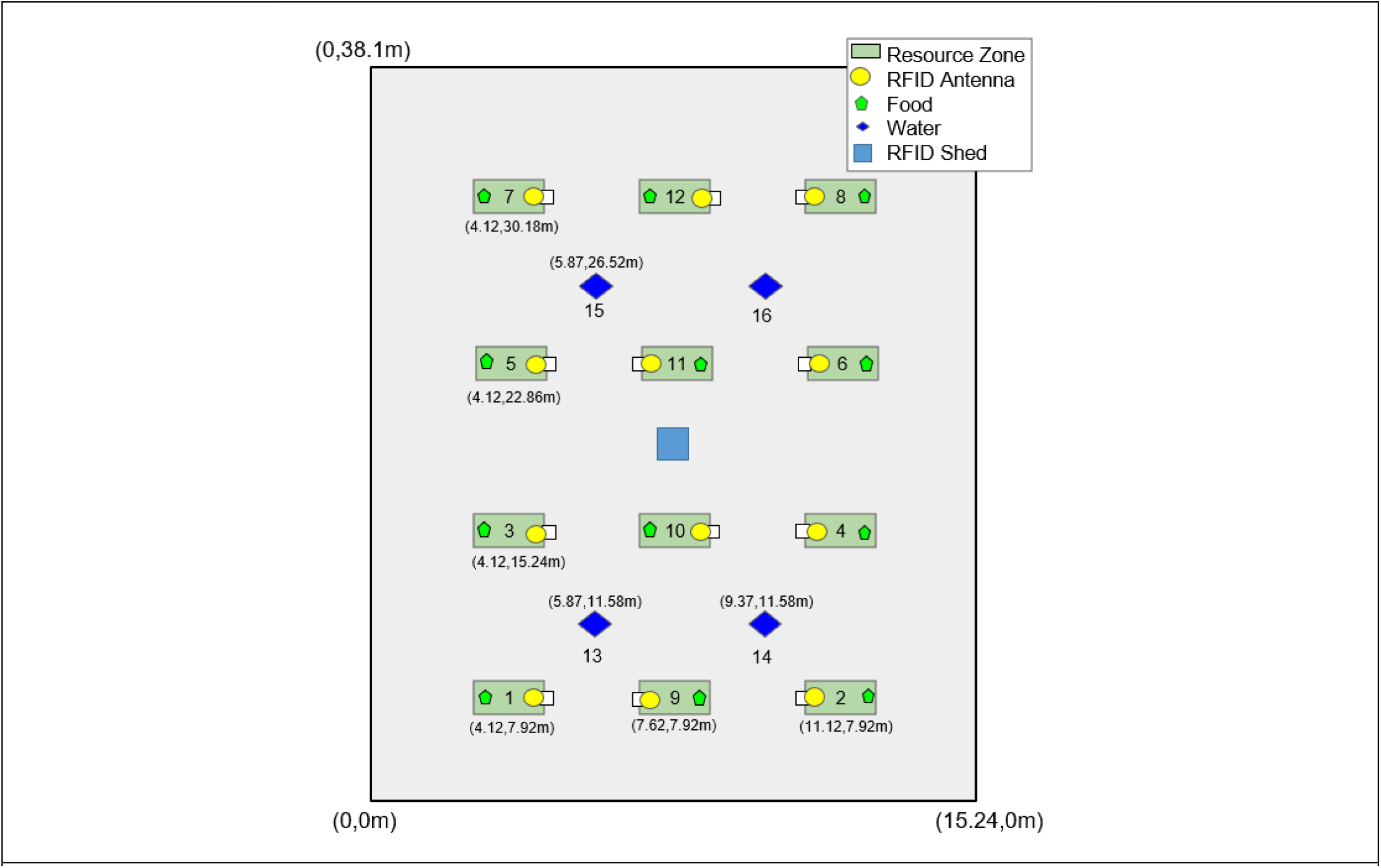
Schematic of the field enclosure and resource zone schematic. 12 resource zones were placed equally spaced apart in a 3×4 grid and contained an RFID antenna and food ad libitum. White boxes indicate the PVC entrance tubes on each of the resource zones. Coordinate measurements are from the center of the resource zone and water towers. Note that the image is not to scale.

**Figure S2.**
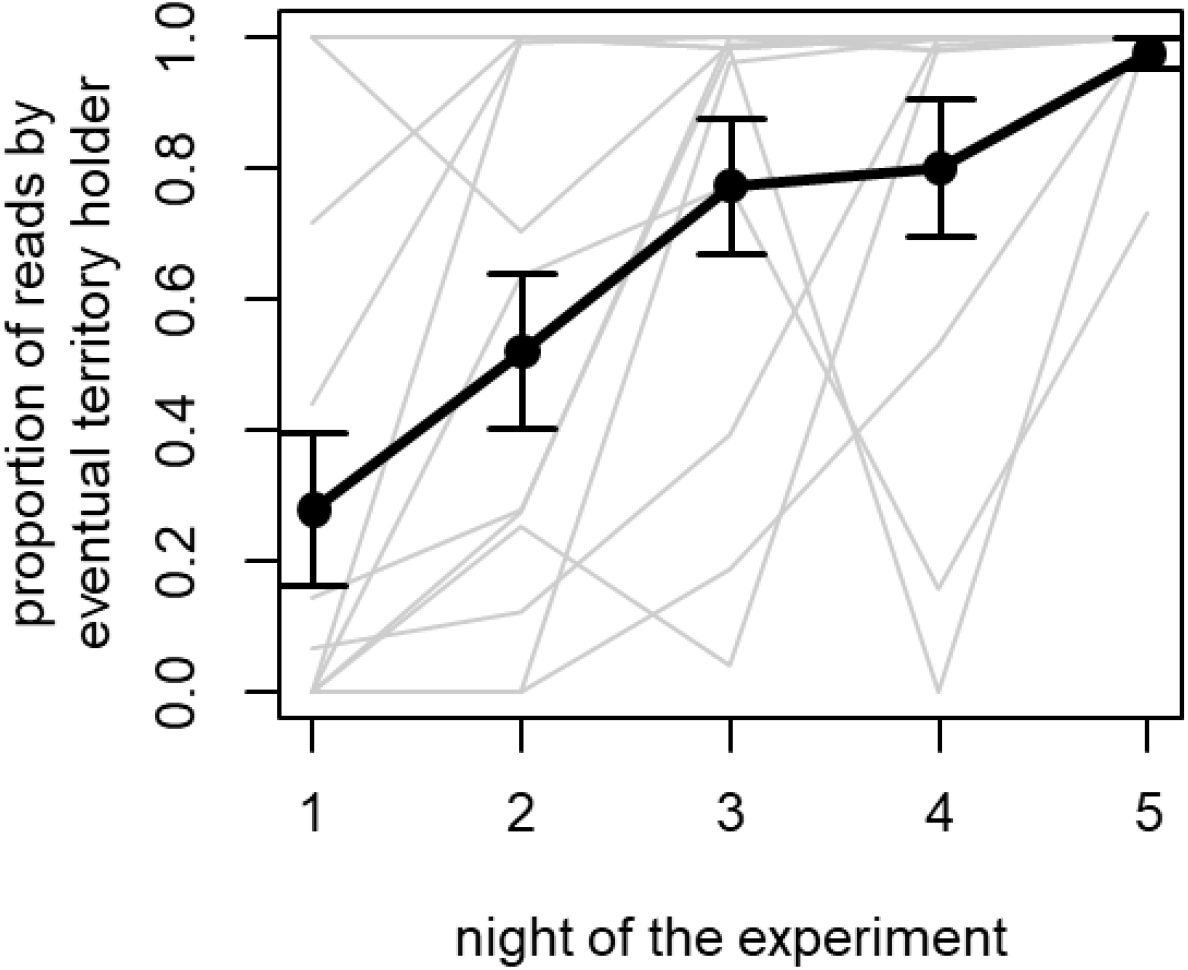
Males acquired territories during the first five nights of the experiment. Gray lines indicate changes in the proportions of RFID reads in each resource zone (n = 12) that originated from the eventual territory holder. The dark points and connecting line represent the mean of these 12 individual measures, with error bars indicating standard error. The territory holder of a given zone was identified as the male that was responsible for the most RFID reads in that zone on night 5 of the experiment. On night 5, nearly all (99.97%) RFID reads that were recorded across all 12 zones originated from the males that controlled them.

**Figure S3.**
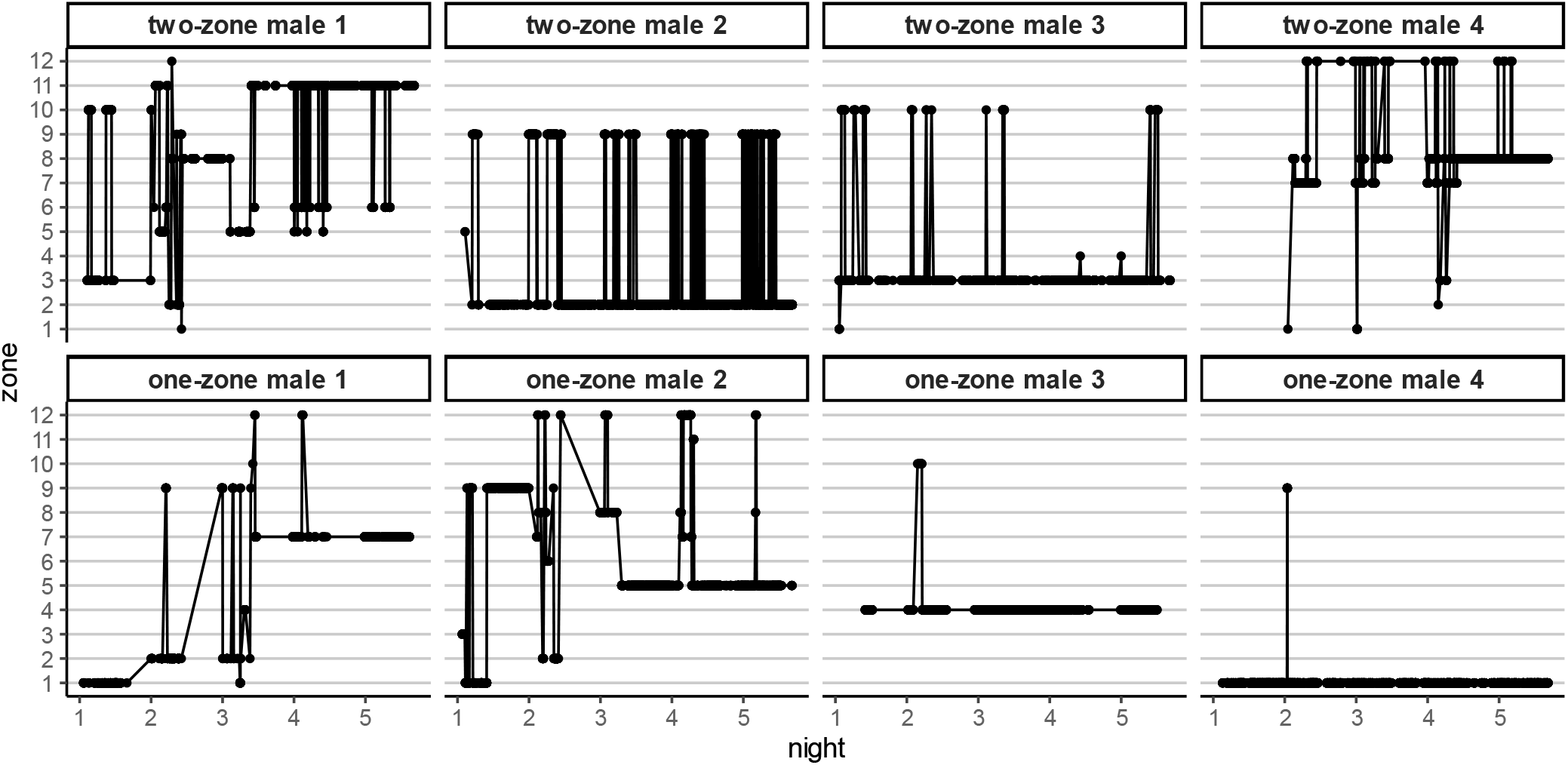
Movement of the 8 original territorial males during their first 5 nights (x-axis) in the enclosure. Each point indicates an RFID read and each line between points indicates transitions between zones (zone locations indicated on y-axis). Note that the four animals in the top row eventually established territorial control over two resource zones between which they made regular nightly visits. The four animals in the lower row established territorial control over one zone each. The addition of intruder males occurred after these territories were established (night 6, not pictured).

**Figure S4.**
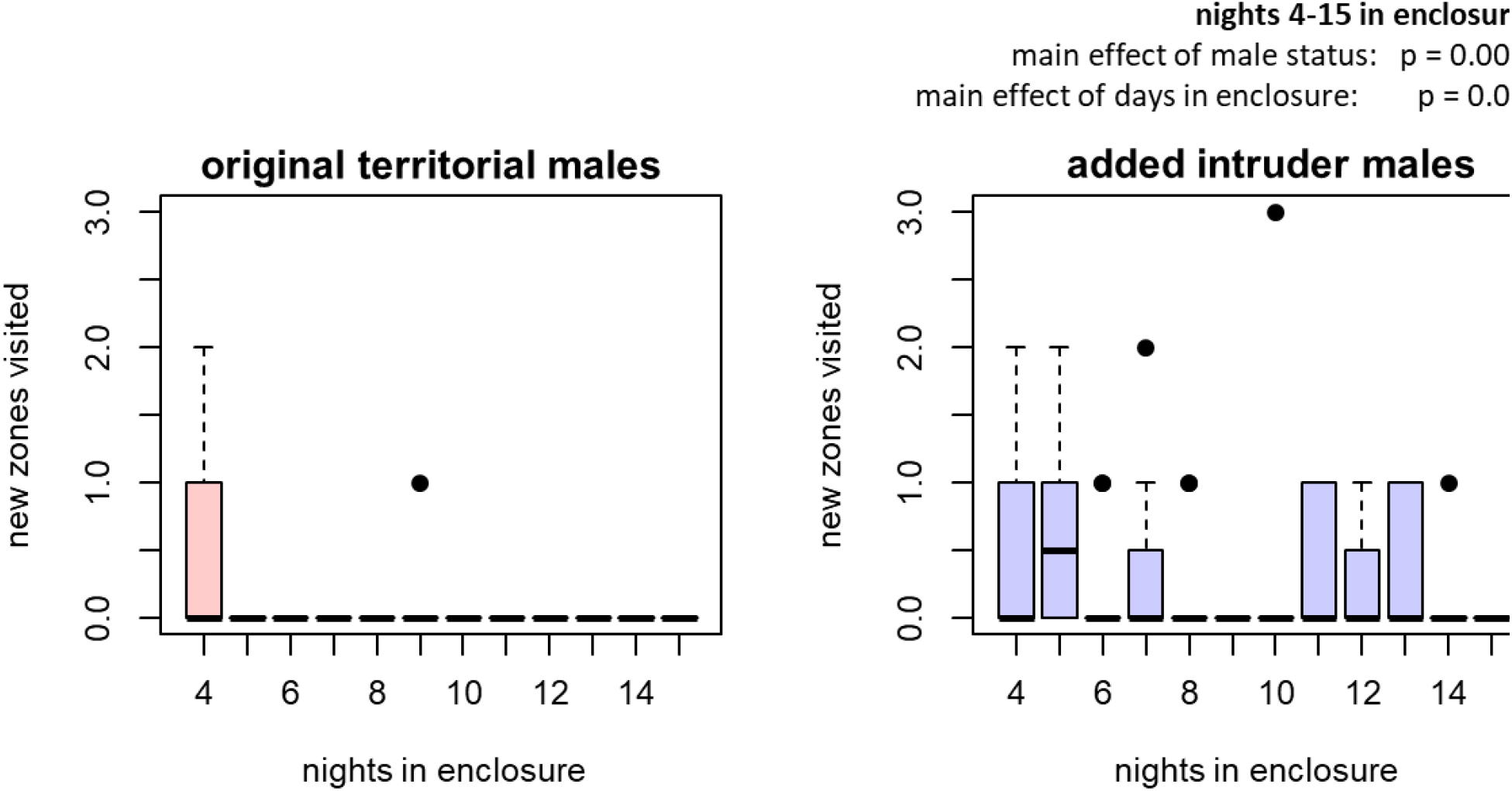
Male exploratory behavior differed, depending on whether the male encountered an environment without any occupied territories (left) or an environment with all territories already filled (right). The y-axis represents the number of new zones that a male visited each night that he had not previously visited. This figure represents a comparable analysis to that presented in Figure 4 of the main text, with the y axis measuring the nightly slope of curves in that figure. As in the main text, added intruder males continue to visit new zones after their third night in the enclosure, as measured in the main effect of this figure and the interaction term in Figure 4.

